# Characterization and Applications of Evoked Responses During Epidural Electrical Stimulation

**DOI:** 10.1101/2023.01.10.523453

**Authors:** Nishant Verma, Ben Romanauski, Danny Lam, Luis Lujan, Stephan Blanz, Kip Ludwig, Scott Lempka, Bruce Knudson, Yuichiro Nishiyama, Jian Hao, Hyun-Joo Park, Erika Ross, Igor Lavrov, Mingming Zhang

## Abstract

Epidural electrical stimulation (EES) of the spinal cord has been FDA approved and used therapeutically for decades. However, there is still not a clear understanding of the local neural substrates and consequently the mechanism of action responsible for the therapeutic effects. Epidural spinal recordings (ESR) are collected from the electrodes placed in the epidural space. ESR contains multi-modality signal components such as the evoked neural response (due to tonic or BurstDR^™^ waveforms), evoked muscle response, stimulation artifact, and cardiac response. The tonic stimulation evoked compound action potential (ECAP) is one of the components in ESR and has been proposed recently to measure local neural activation during EES. Here, we first review and investigate the referencing strategies, as they apply to ECAP component in ESR in the domestic swine animal model. We then examine how ECAP component can be used to sense lead migration, an adverse outcome following lead placement that can reduce therapeutic efficacy. Lastly, we show and isolate concurrent activation of local back and leg muscles during EES, demonstrating that the ESR obtained from the recording contacts contain both ECAP and EMG components. These findings may further guide the implementation of recording and reference contacts in an implantable EES system and provide preliminary evidence for the utility of ECAP component in ESR to detect lead migration. We expect these results to facilitate future development of EES methodology and implementation of use of different components in ESR to improve EES therapy.

## Introduction

Epidural electrical stimulation (EES) has been used for several decades to manage chronic pain (Shealy et al., 1967)(Melzack & Wall, 1965). Despite this clinical adoption, the local neural substrates responsible for the therapeutic effects of EES, and consequently the mechanisms of action to control pain, are poorly understood. Activation of the large-diameter fibers in the dorsal columns is thought to inhibit pain transmission via gating within the dorsal horn (Zhang et al., 2014) (Yang et al., 2011) however, several studies suggest that additional segmental and supraspinal mechanisms are involved in EES-induced analgesia (Gilbert et al., 2022). Variations in the effect of EES to control pain can be attributed to multiple factors, such as initial placement or later migration of the electrode contacts, patient position, and the functional state of the neuronal circuitry (Mekhail et al., 2022) (Pahapill et al., 2020) (Dombovy-Johnson et al., 2022a). Although lead migration can be confirmed through clinical imaging, there is currently no clinically available method to detect and flag the possibility of lead migration for EES automatically and continuously which would allow for decreased loss of efficacy.

Several approaches have been developed to optimize the electrode placement and stimulation parameter selection of EES with the goal of reducing variation in and increasing therapeutic effectiveness of EES. These range from the classical mapping of paresthesia during trial stimulation to recently introduced closed-loop control based on motion/position sensors and/or evoked compound action potential (ECAP) sensing following stimulation. The long-term decline of EES efficacy represents another significant problem leading to the explant of stimulation devices. (Pope et al., 2017).

Our other studies about epidural spinal recordings (ESR) showed that ESR, collected from electrodes implanted in the epidural space, are comprehensive recordings containing multimodality signals, such as tonic stimulation evoked responses (ECAP component), BurstDR^™^ stimulation evoked charge accumulative evoked responses, stimulation evoked electromyogram responses (EMG component), cardiac signals. Real-time analysis of ECAP component and EMG component in ESR during EES may help to optimize EES protocols by enabling real-time adjustment of stimulation contacts and parameters. Moreover, the ECAP component in ESR may provide a clinically viable method to automatically detect lead migration if morphology of the ECAP components changes with migration of the recording or stimulation lead (Calvert et al., 2022). The work by Calvert and colleagues (2022) investigated the migration detection application with paddle leads, which require a laminectomy to place and were designed specifically for the restoration of motor function. Paddle leads are also less common in the treatment of chronic pain than the percutaneously placed cylindrical leads. The different physiological components, together with its derived features, could be also measured in real-time to provide feedback on neural substrates activated during EES (Melzack & Wall, 1965)(Russo et al., 2018).

This study is focused on 1) understanding the fundamentals of electrophysiology recordings, particularly referencing strategy, to ESR during EES; 2) investigating changes in ECAP component morphology during shifts of the stimulation and recording leads to explore lead migration detection as an application of ECAP component in ESR; 3) investigating the mechanisim of activation of local back muscles during stimulation and the detection of EMG component in ESR by the implanted leads. In all cases, we use controls, including recordings made in the animal after euthanasia (no evoked neural response after death), to differentiate bona-fide ECAP components from stimulation artifact and ringing induced by filtering of stimulation artifact (Nicolai et al., 2020).

## Methods

### Subjects

All study procedures were conducted with the approval of the Mayo Clinic Institutional Animal Care and Use Committee and in accordance with the National Institutes of Health Guidelines for Animal Research (Guide for the Care and Use of Laboratory Animals). Domestic white swine males and females aged 8-12 weeks and weighing 27-46 Kg were used for this study. Animals were kept in separate cages in a controlled environment (constant temperature at 21°C and humidity at 45%) on a 12-hour light/dark cycle with ad libitum access to water and were fed once daily.

### Stimulation and Recording

We performed *in vivo* electrophysiological experiments on all animals to record the ESR from the spinal cord and spinally evoked motor potentials (SEMP) from selected muscles during EES. The surgical approach has been described in detail previously (Cuellar et al., 2017). Briefly, intramuscular telazol (5mg/kg) and xylazine (2 mg/kg) were administered for anesthesia induction and 1.5-3% isoflurane for maintenance. Fentanyl was continuously administered during surgery (2-5 mg/kg/h) for analgesia. Laminectomies were performed to expose the lumbosacral spinal cord (Th1-L1). Connective and fat tissue was removed while keeping the dura mater intact. For EES an 8-contact platinum iridium Octrode^™^lead with 1.3 mm diameter, 3 mm contact length, and 4 mm spacing between contacts was used to preform EES (Abbott, Plano, TX). Two leads were placed on the spinal cord with the stimulation lead shifted rostro-caudally and medio-laterally during the experiment and the recording-only lead shifted only rostro-caudally. To keep the leads secured and stable, minimal laminectomies were performed on L1and the 5 thoracic segments rostral, to pass the leads through and check their position. This approach allowed the leads to be visualized with fluoroscopy and manipulated in relationship to the dorsal spinal cord anatomy, accurately adjusting their position. For the reference electrode, a stainless-steel needle electrode (AS 631, Cooner wire) was inserted in the paravertebral muscles on the left side of the surgical site or contact 3 or 9 on the implanted leads (Fig 1B) was assigned. Four different referencing methods were derived from this set up: (i) Local Tissue Referencing (LTR), which used the needle electrode as the contact; (ii) Differential (DIFF) utilizing subtraction of the more caudal of two neighboring channels; (iii) utilizing the contact furthest contact from the stimulation contacts, thus referencing contact 9 (REF9); and (iv) using contact 3 as reference (REF3). In order to reduce the impact of the stimulation artifact on the ESR the stimulation waveform used was an asymmetric, charge-balanced waveform with an anodic-leading rectangular pulse with a duration of 400 μs followed by a cathodic pulse with a duration of 80μs. The cathodic phase of the waveform had an amplitude five times greater than the amplitude of the leading anodic phase (Grill & Mortimer, 1995). Stimulation amplitude was defined as the amplitude of the cathodic phase. The contacts used for stimulation were contacts 7 and 8 (Fig 1). The stimulation waveform was delivered at 38 Hz with the battery isolated Subject Interface Module (SIM) from Tucker Davis Technologies (TDT). Further, the TDT WS4 Computer, RZ5D Bioamp Processor, and the IZ10 Stimulation/amplifier, were used in conjunction with Synapse software for stimulation and recording.

**Figure 1:**
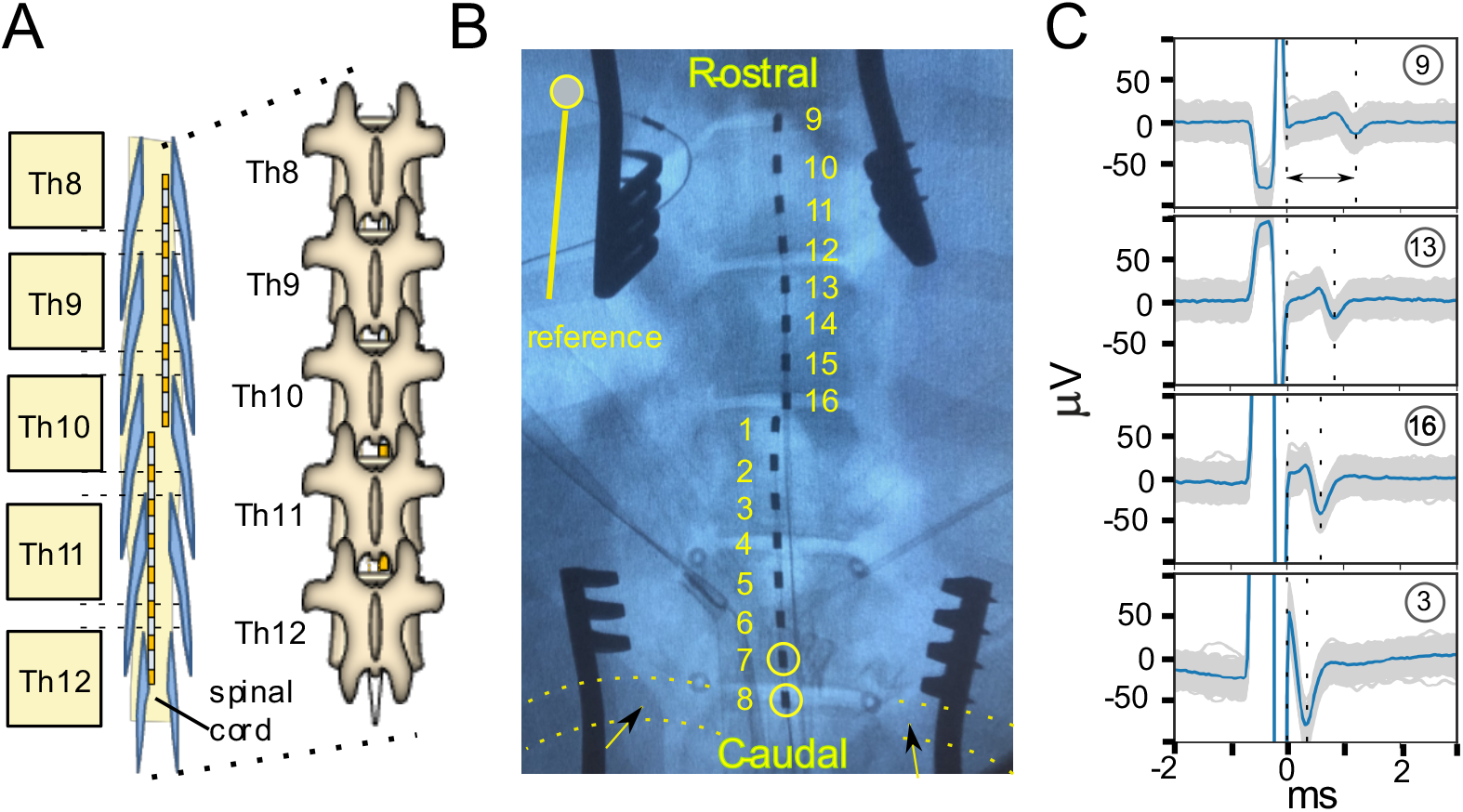
Lead placement on spinal cord and electrophysiology recording examples. (A) cartoon demonstration of two Octrode^™^ leads placed in series on the spinal cord with partial laminectomies starting on L1 and the 5 thoracic segments rostral which can be seen in the animated spinal column. (B) X-ray image showing the placement of two leads with the lowest contact (8) located at the same level as the lowest rib bone (marked with arrow and outlined with dotted lines). Electrode contracts 7 and 8 on the first lead were used as stimulation electrodes, unless otherwise stated. The second lead with contacts 9 to 16 was placed more rostrally. (C) Example of ESR from four different channels along the electrode arrays. In each subplot, the grey background is the aggregation of 300 individual recordings, and the blue trace is the median waveform of these consecutive individual trials. The two dashed lines in each subplot indicate the end of stimulation and the biggest negative peak in the ECAP component in ESR, respectively. The arrowed line in the top subplot shows the measured latency of the ECAP component recorded on Channel 9.

### ESR and SEMP recordings

ESR were recorded concurrently on all 14 non-stimulation electrodes along the two implanted Octrode^™^ leads (Fig 1B). The two designated stimulation contacts, 7 and 8, are circled in Fig 1B. For recording SEMPs, a pair of two stainless steel needle electrodes were placed intramuscularly into the two lowest intercostal muscles (IM), the rostral portion of the paravertebral muscles (PVR), the caudal section of the paravertebral muscles (PVC), and into selected muscles of the hind limbs (i.e., gluteus (G), quadriceps (Q), and bicep femoris (BF)). The signals were recorded and digitized (sampled at ~25 kHz). Offline, the recorded responses were band-pass filtered (100 Hz high-pass, first-order Butterworth and 3 kHz low-pass, Gaussian) in Python 3.7 with the SciPy package.

### Data and statistical analysis

The pyeCAP package (https://pypi.org/project/pyeCAP/) was used in Python 3.7 for the offline handling and analysis of electrophysiology data. For the analysis of the impact of referencing method on stimulation artifact and ECAP component amplitude, stimulation artifact magnitude was quantified as the peak-to-peak signal of the averaged recording (median of 150-300 traces). ECAP component amplitude was quantified as the peak-to-peak signal of the averaged ESR trace between 2.1 ms and 5 ms, which is the duration immediately following the stimulation artifact while encompassing the entire ECAP signal. Note that stimulation was only delivered starting at ~1.3 ms. For the analysis of referencing method on stimulation artifact and ECAP component amplitude, stimulation artifact and ECAP component amplitude for LTR reference was normalized to 1 for each subject to account for subject-to-subject variability and enable cohort-based analysis. Shapiro-Wilk’s test was used to test the samples from n=5 subjects for normality to satisfy the assumption of the subsequently used one sample or paired t-test (twotailed *α* = 0.05).

To determine the onset and amplitude of each SEMP, waveforms were analyzed and compared starting from the current threshold that elicited the onset of SEMP. SMEP peak-to-peak response amplitudes and latencies were measured in a window of 5 to 18 ms following the stimulation artifact. One-way ANOVA with post-hoc Tukey’s Honestly Significant Difference (HSD) test was used to test for changes in ECAP latency and amplitude from different electrodes.

This study was not pre-registered and so all findings should be considering exploratory and not confirmatory.

## Results

### Sample ECAP components in ESR recording and quantification

Figure 1C shows ESR recordings from selected electrode contacts 3, 16, 13, and 9 along the caudal-rostral axis while stimulation was applied from contacts 7 (anode) and 8 (cathode) on the caudal lead. This example shows that at a stimulation amplitude of 1.0 mA, the stimulation artifact and ECAP component could be recorded in ESR along the caudal-rostral axis from the selected contacts (Fig 1C). Within each channel, the ECAP latency was defined as the time between the end of the stimulus artifact to the highest negative peak of the ECAP component as illustrated in Fig 1C (top). From contact 3 to contact 9, the measured latency was increased from 0.35 to 1.22 ms, which indicates the propagation of action potentials in the rostral direction. The ECAP amplitude decreased as the recording electrodes located further from the stimulating electrodes. From contact 3 to contact 9, the measured amplitude decreased from 103 to 19 □ V.

### Effect of reference electrode location on morphology of stimulation artifact and ECAP

The first question explored in this study is the role of referencing in ECAP component in ESR. Differential reference (DIFF) was compared with local tissue reference (LTR) and reference on contact 9 (REF9), which is the recording contact furthest from the stimulation contacts. Fig 2 demonstrates representative variations in ESR and stimulation artifact magnitude and morphology of ECAP component with the different referencing methods. The data showed that recordings using DIFF had the lowest stimulation artifact magnitude (0.529 ± 0.046 standard error (SE)), while recording using LTR (normalized to 1 in each of n=5 subjects) and REF9 (0.998 ± 0.051 SE) had higher stimulation artifact magnitudes (P<0.001 for DIFF compared to LTR and REF9) (Fig 2A and 2C). The reduced stimulation artifact for the DIFF reference was due to the proximity of the reference electrode to the recording electrode and hence the similar representation of the stimulation artifact on both the recording and reference electrode, which subtract from one another. Note the distance between the DIFF recording and reference pair affects how similar the representation of the signal is on both electrodes and how pronounced this subtraction effect is. Similarly, ECAP component magnitude was significantly higher with LTR (normalized to 1 in each of n=5 subjects) and with REF9 (1.029 ± 0.051 SE) compared to the DIFF referenced recordings (0.737 ± 0.069 SE) (P<0.005 for DIFF compared to LTR) (Fig 2A and 2D).

**Figure 2.**
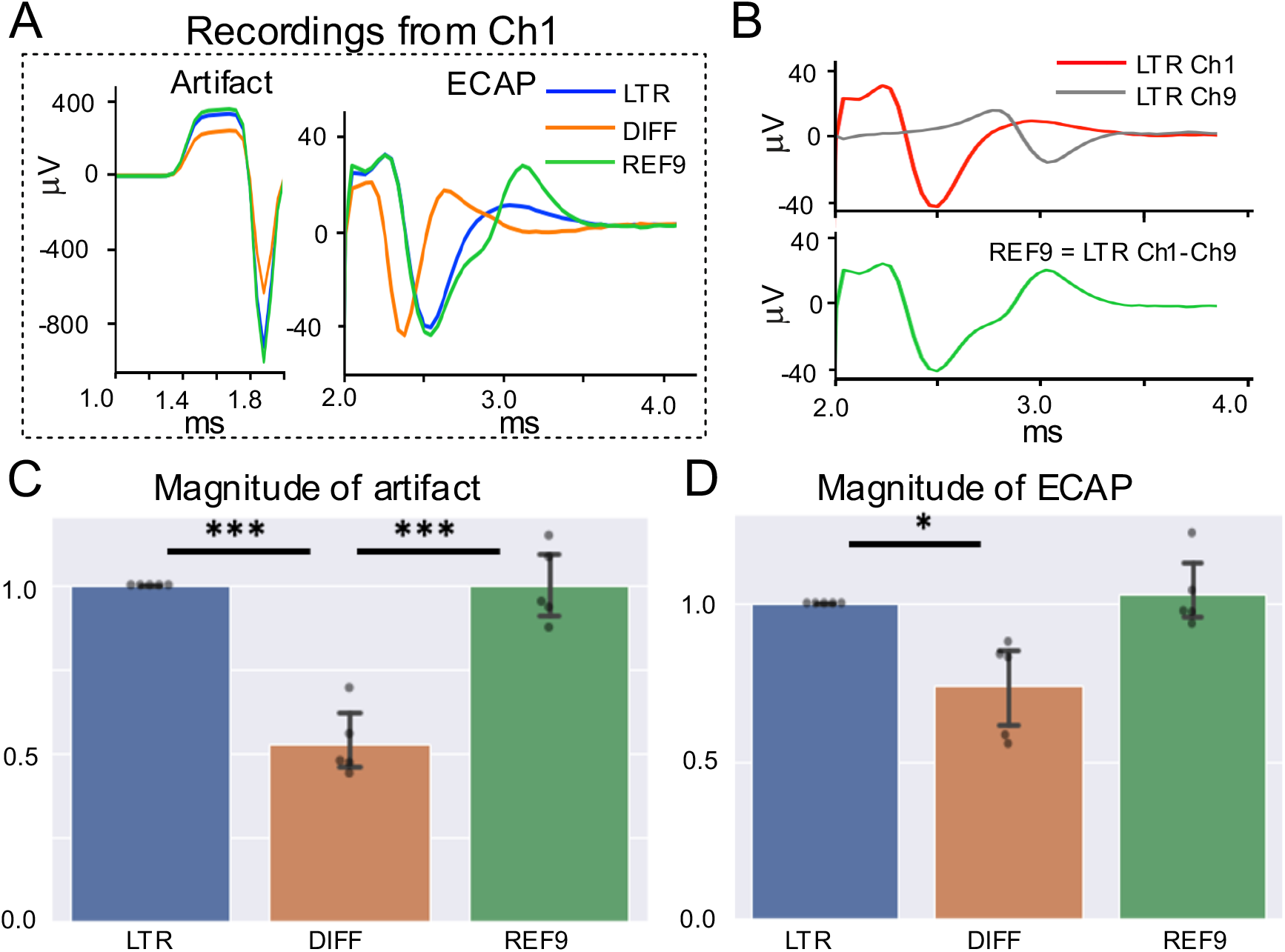
Effect of reference electrode placement on stimulation artifact and ECAP morphology in ESR. (A) Representative plot of stimulation artifact (left) and ECAP component (right) recorded on channel 1 (Ch1) using different referencing options: local tissue reference (LTR), differential recording reference using channel 2 and channel 1 (DIFF), and on lead reference using contact 9 (REF9). (B) Experimental data illustrating how a recording using REF9 can be computed from channel 9 (Ch9) using LTR ‘subtracting’ into the recording from Ch1 using LTR. The waveform in each plot is the median waveform from 150-300 number of trials with stimulation amplitude at 2.4 mA. (C) Magnitude (peak-to-peak) of the stimulation artifact (n=5 pigs). (D) Magnitude (peak-to-peak) of the ECAP component (n=5 pigs). * Indicates significant difference in the ECAP magnitude (p<0.05) and *** indicates (p<0.001). Error bars represent one standard deviation.

The ESR trace recorded using REF9 has an additional inflection in its morphology due to the ‘subtraction’ of the same ECAP component recorded concurrently on the reference and the recording electrode (Ch1) (Fig 2B).

These data suggests that LTR recordings provide the clearest representation of the neural activity (no ECAP subtraction effects). However, is prone to common mode artifact contamination (e.g., stimulation artifact) and requires an additional electrode in non-neural tissue. LTR could be implemented clinically with reference to the metal external of the implantable electronics. DIFF reference was most robust to common mode artifact but also resulted in diminished neural signal magnitude. REF9 provided a middle ground with ease to implement and a clear representation of the neural activity with minimal ECAP component subtraction effects from the reference electrode. For the rest of this study, LTR was used as it provided the best representation of underlying neural activity and could be virtually re-referenced to both REF9 and DIFF.

### Variation of ECAP amplitude and latency in ESR across recording channels with different reference options

Fig 3A shows that the ECAP magnitude changed in different ESR as it conducted from the recording contacts closer to the stimulation contacts to the recording contacts further from the stimulation contacts. The role of referencing method on conduction latency across recording channels is demonstrated in Fig 3B with concurrent ESR from all available channels. The difference in conduction delay across the channels is critical to consider for measuring the conduction velocity of an ECAP component and verify its authenticity from common mode artifact, which appears simultaneously on all contacts. Recordings using LTR and DIFF reference demonstrate a more linear change in the ECAP latency across the leads indicating approximately constant conduction delay between contacts (Fig 3B, first and second from the left). Recordings using REF9 reference shows a similar pattern in the initial recording contacts located more caudally (closest to stimulation contacts). However, on recording contacts closer to the reference contact 9 located most rostrally, the conduction delay between subsequent contacts diminished (Fig 3B, third from the left). This may be because the ESR is recorded on both the active recording contact and the reference electrode contact 9 (Fig 2B). Lastly, contact 3 was used as the reference electrode (REF3), and the morphology of the ECAP component was unique compared to other three referencing methods. Particularly, the latency delay predictably decreased as the channels were moving more rostrally from the physical location of contact 3 (Fig 3B, fourth from the left).This may be because the ECAP is being recorded at the same time on contact 3 for each recorded channel resulting in a similar effect that is seen in REF9 (Fig 2B) and could results in the polarity flip seen below and above contact 3.

**Figure 3.**
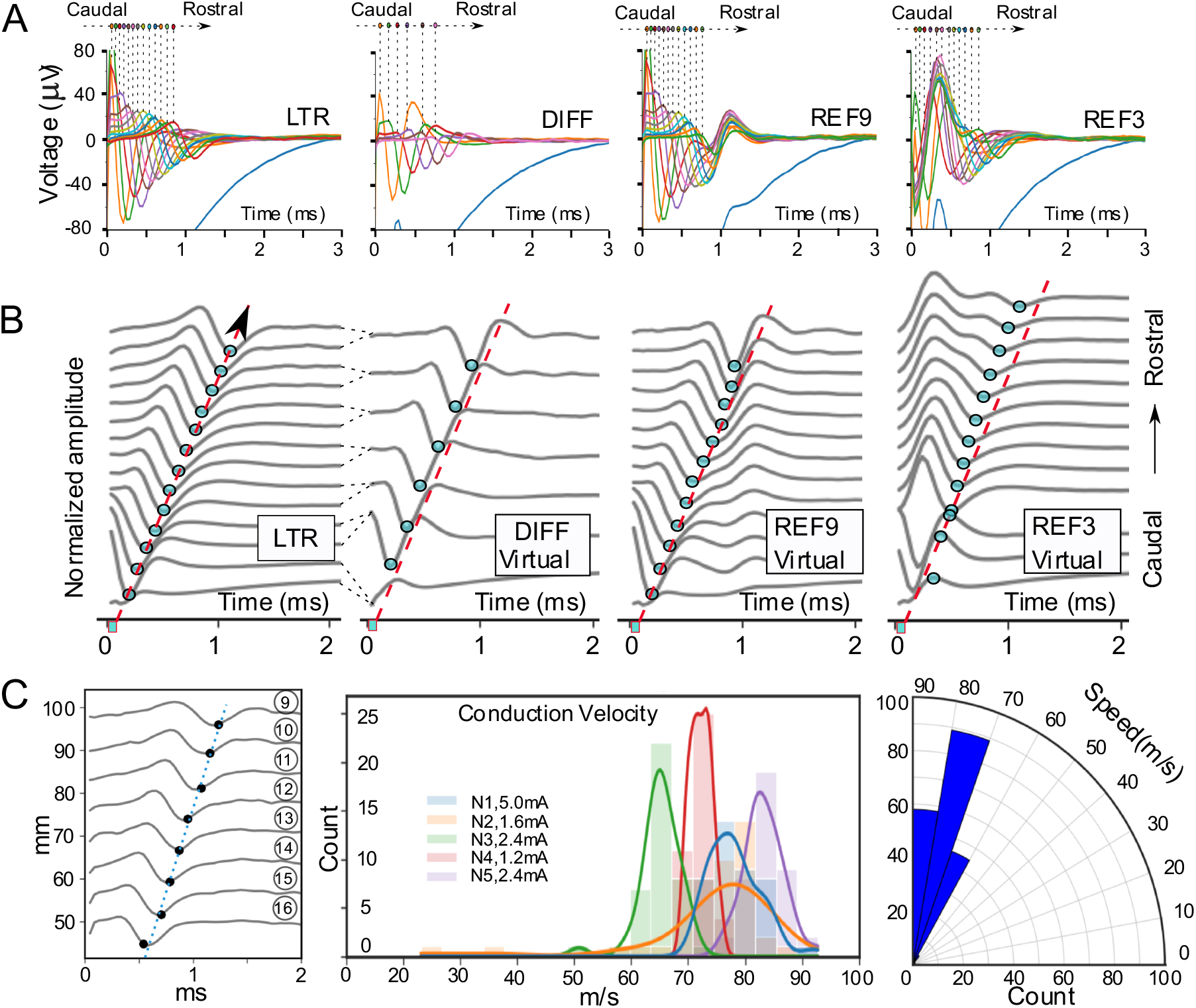
The effect of referencing method on ECAP component amplitude and latency. (A) Representative recordings on all available recording channels demonstrate changes in ECAP magnitude with local tissue reference (LTR), differential reference (DIFF), contact 9 as reference (REF9), and contact 3 as reference (REF3). Each recording trace is a median waveform of 150 trials for a stimulation amplitude of 2.4 mA. (B) The influence of referencing method on conduction latency across lead recording channels. ECAP component in ESR made simultaneously across all available recording channels on the leads from caudal to rostral location (refer to Fig 1 for more information about electrode position) with different referencing (LTR, DIFF, REF9, REF3) done virtually. Each recording trace was a median waveform of 150 trials and was normalized to itself. Dots on each trace indicate the negative peak in each channel. The dots in the first subplot using LTR are traced with a red dashed line indicating the propagation of the ECAP. The red dashed line in the following subplots are at identical slant angle and position to the one in the first subplot using LTR reference. (C) (Left) Illustrates ECAP propagation from channels 9 to 16 on the rostral recording Octrode^™^ lead. Negative ECAP peak location was detected and marked in each channel for latency measurements. Conduction velocity was estimated using a linear regression between the recording-stimulating contact distances and ECAP latency (as indicated by the dotted line). (Middle) ECAP conduction velocities were extracted and plotted at different stimulation amplitudes (1.2 to 5 mA) from five different subjects. For each subject, the density plot is shown overlayed onto a histogram. Most of the measured ECAP conduction velocities fell within the range for Aβ-fiber activation, 35-80 m/s. (Right) Radar plot shows the projection of ECAP conduction velocities with data from all subjects grouped.

### Quantification of conduction velocity

Quantifying the conduction velocity of the ECAP component is useful to determine the underlying fiber types recruited. The Aβ-fiber type is known to have a conduction velocity in the range of 35-80 m/s (Bear et al., 2007), Aα-fibers in the range of 80-120 m/s, and Aδ-fibers in the range of 5-35 m/s. The recordings collected from the recording Octrode^™^ lead was used to calculate the ECAP conduction velocity. Recordings from the recording Octrode^™^ were used to minimize the distortion caused by the stimulation artifact prevalent on the recording contacts on the stimulation lead. Due to the variability in stimulation thresholds between subjects, stimulation amplitudes were selected where the ECAP was clearly observed but under the motor thresholds for that subject. A single dataset contained 300 trials of stimulations. To ensure consistencies in calculating ECAP propagation, the median wave of every 5 trials was obtained for each channel. In total, 40 median trials per subject were extracted for final conduction velocity calculations, excluding trials that were not consistent throughout Channels 16 to 9. A linear regression model was developed using the temporal locations of the negative peaks of ECAP signals in ESR from recording channels and their respective contact distances from the stimulating contact (Fig 3C, first from the left). An example of ECAP propagation in Figure 3C (left) on the recording lead was shown to have a conduction velocity of 72.29 m/s, at the boundary of the A *β* -fiber and A *α* -fiber conduction velocity ranges. Fig 3C (second from the left) summarizes the conduction velocity results across subjects. Stimulation amplitudes across the five subjects fell within the range of 1.2 – 5.0 mA, with no conduction velocity distribution pattern apparent with stimulation level across subjects.

### ECAP component variations in ESR show recording lead migration

Lead migration is a common complication that leads to reduced long-term efficacy of EES (Dombovy-Johnson et al., 2022b). The ECAP component in ESR could be used to continuously sense and flag the possibility of lead migration. In this study we investigate the effects of rostro-caudal and medio-lateral lead migration on ECAP morphology, area-under-the-curve (AUC), and latency as features that could be implemented to detect lead migration. The caudal Octrode^™^ lead was used for stimulation and was fixed while the rostral lead was shifted to record signals at multiple locations (3.5, 28, and 52.5 mm from the initial arrangement *‘I’*) (Fig 4A). Stimulation-recording distance was defined as the distance between the stimulation contacts to the selected recording contact. The latency for each ECAP component was defined as the time delay between the end of stimulation to the maximum negative peak in the ECAP signal, while the strength of ECAP signal is defined as the AUC of the ECAP window, which is indicated by the two dashed lines in ESR shown in Fig 4B. Fig 4C shows representative changes in ECAP latency and strength at four different locations (IIV). Next, datasets from tested animals were selected for stimulation amplitudes ranging from 2 to 2.5 mA at which a clear ECAP signal was visually observed but without contamination from a strong EMG signal caused by an excessive stimulation dose. As the recording lead was shifted rostrally, the ECAP latency increased as the stimulation-recording distance increased. The averaged latency obtained from recording at each of the four locations (*I-IV*) was 0.462 ± 0.117 ms, 0.598 ± 0.171 ms, 0.890 ± 0.136 ms, and 1.145 ± 0.155 ms, respectively (Fig 4D). Comparing latency measured at the baseline position *‘I’* to that obtained from other positions *‘II’, ‘III’, and ‘IV’*, the increases were all significant (p<0.001). ECAP component strength, as measured by AUC, were normalized to respective stimulation amplitudes. For each rostral shift, ECAP strength was measured as percent change from the baseline position (Fig 4E). When compared to the baseline, ECAP AUC decreased as the recording lead was moved rostrally. The changes were - 24.1 ± 13.1% (from position *I* to *II*), −67.7 ± 5.6% (from position *I* to *III*), and −85.6 ± 11.0% (from position *I* to *IV*). As the distance between the stimulation and recording contacts increased, there was a larger reduction in ECAP strength, and the change was significant across all groups (p<0.001). These data show significant changes in ECAP latency and strength with lead migration, suggesting that the migration of one lead relative to the second implanted lead could be detected by changes in ECAP morphology.

**Figure 4:**
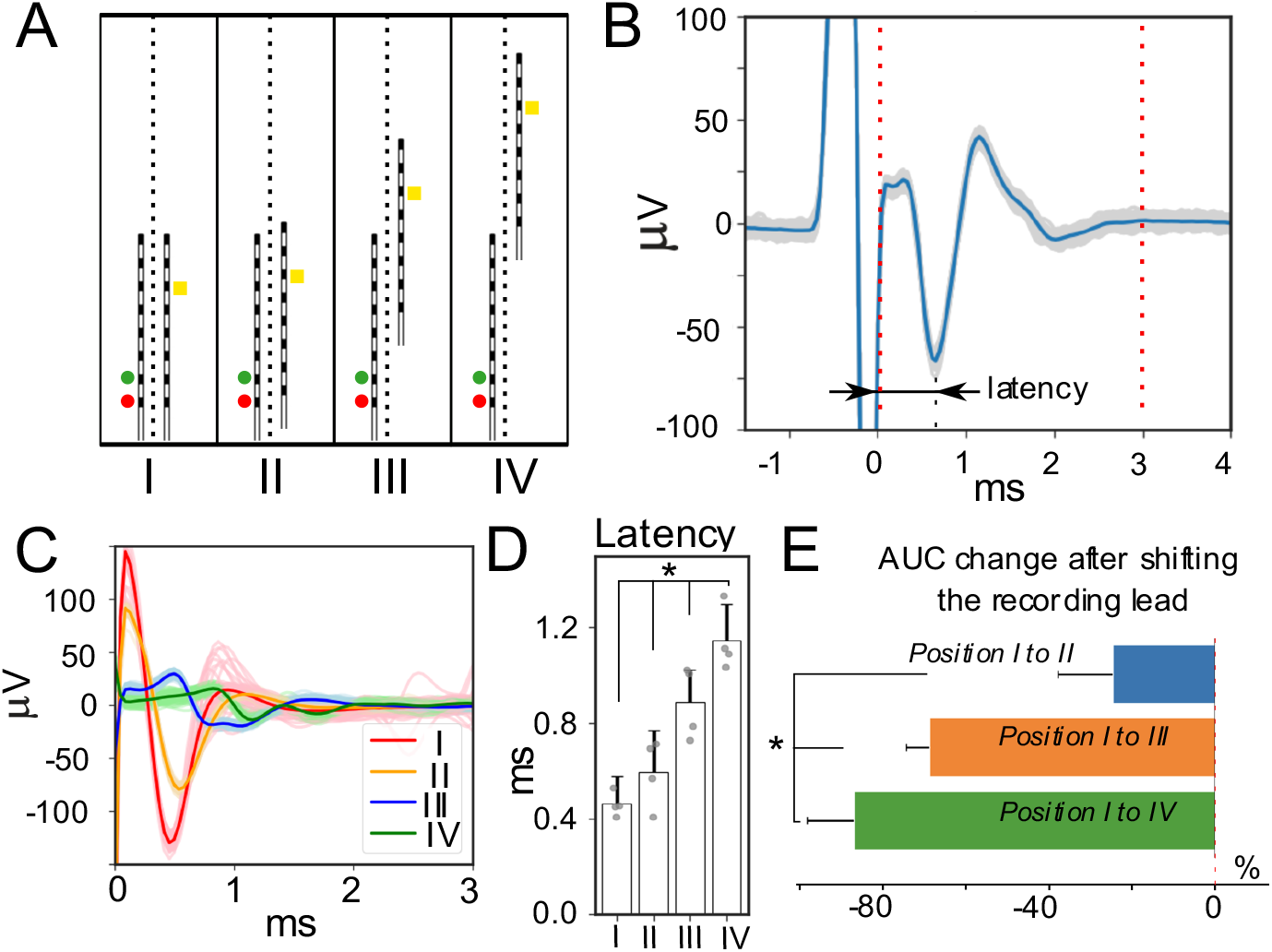
Recording lead migration affects features of the ECAP component in ESR. (A) X-ray images of implanted leads with recording lead shifting rostrally to three predetermined locations. Circular markers located left of the first lead indicate stimulating contacts 7 (yellow, anodic) and 8 (red, cathodic). Contact 11 (white square) located on the second lead indicates the recording channel that was used for the reported ECAP component quantifications. The four different recording locations were defined as location *‘I’, ‘II’, ‘III’, and ‘IV’*. (B) Median representative waveform out of 300 recorded trials with the ESR following the end of the stimulation artifact. Latency was measured as the time between the end of the stimulation artifact and the maximum negative ECAP component peak. (C) Median waveform from 300 trials recorded in an animal at all four locations with a stimulation amplitude of 2 mA. Here, 0 ms indicates the end of stimulation. (D) Averaged latency across subjects (n=4) showing the latency increased as the recording lead was shifted rostrally. ECAP latency increased significantly as the recording electrode was shifted rostrally for all lead configurations (p<0.001). (E) Percent changes to ECAP strength (shown as area-under-the-curve, AUC) are plotted averaged across all subjects (n=4). As the distance between the stimulating and recording contacts increased, the percent decrease in the ECAP magnitude compared to the magnitude at the initial position, *I*, increased (p<0.001 tested against null hypothesis of no change in ECAP magnitude).

### The effect of lead migration along rostral-caudal and medial-lateral axis on the ECAP component morphology

Next, we investigated if rostro-caudal or medio-lateral shifts in the lead with the stimulation contacts could be detected by the second recording lead. The stimulation lead was shifted approximately 24.5 mm from a more rostral position to a more caudal position (Fig 5A). All ESR included in this analysis had clearly observed ECAP components. 150 individual stimulation trials were used from each subject. Fig 5B shows a representative trace illustrating that the recorded ESR in the caudal position have a longer latency in ECAP component as the stimulation-recording distance increases after shifting from the rostral location (Fig 5B). Two distinct populations for ECAP component latency are shown in the histogram plot (Fig 5C), showcasing differences pertaining to the respective lead locations. After the caudal shift of the stimulation lead, the ECAP component latency was significantly increased from 0.424 ± 0.024 ms in the rostral position to 0.860 ± 0.077 ms in the caudal position (p<0.001) (Fig 5C). There was also a substantial reduction in ECAP component strength, reported as AUC change for all subjects: N1: 98.7 ± 43.7 %; N2: 277.2 ± 128.9%; N3: −11.0 ± 8.3% (outlier); N4: 532.1 ± 94.7% (p<0.001) (Fig 5D). The reduction in ECAP component strength is likely driven in large part by the increased distance between the stimulation and recording contacts, thus, limiting stimulation artifact addition into the AUC calculation in quantifying ECAP component.

**Figure 5:**
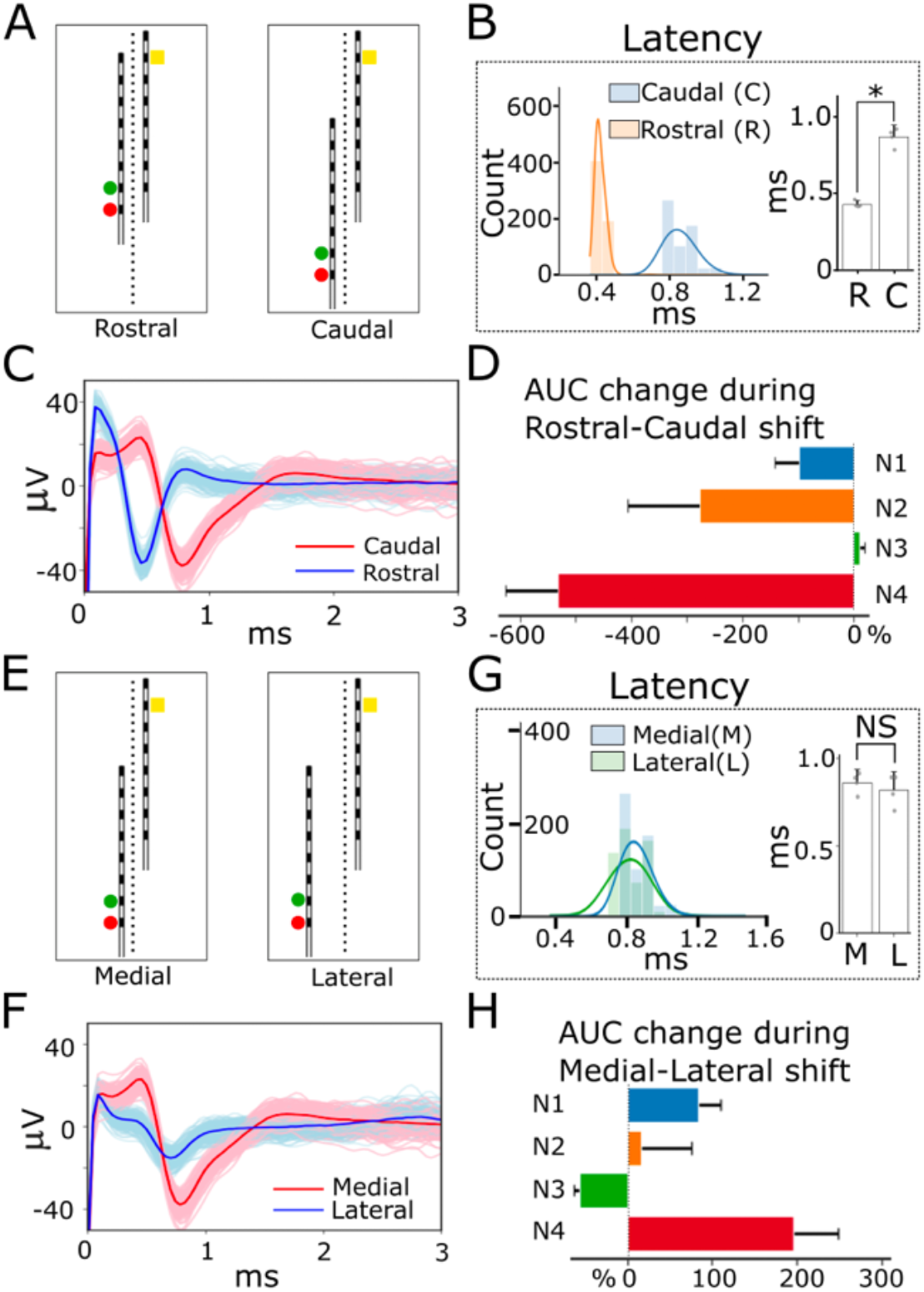
Effects of migration of the stimulating lead along the rostral-caudal and medial-lateral axes on the ECAP component in ESR. (A) X-ray images showcasing the caudal-rostral movement of the stimulation lead. Circular markers located left of the lower lead indicate stimulating contacts 7 (yellow, anodic) and 8 (red, cathodic). Contact 11 (yellow square) located on the second lead indicates the recording channel that was used for the reported ECAP quantifications. (B) ECAP latency calculated for stimulation lead in rostral and caudal position for all subjects (n=4). The histogram shows the distribution of the latency measured at both locations. The bar plot indicates there was a significant increase (p<0.001) in latency as the stimulation-recording distance increased. (C) Example median ESR waveforms from channel 11 are plotted overlain on 300 raw traces at 4.8 mA of stimulation for both the caudal and rostral position revealing changes in ECAP component latency. (D) From rostral to caudal shift in the stimulating lead, the normalized ECAP magnitude (AUC) decreased for all but one subject. (E) X-ray images illustrate stimulation lead shifted more laterally. Circular markers located left of the lower lead indicate stimulating contacts 7 (yellow, anodic) and 8 (red, cathodic). Contact 11 (yellow square) located on the second lead indicates the recording channel that was used for the reported ECAP quantifications. (F) Example median ESR waveforms are plotted overlain on 300 raw traces at 80% motor threshold for both medial and lateral configurations from channel 11. (G) ECAP component latencies were compared for medial and lateral electrode positions (n=4 subjects). Shifting the stimulation lead along the medial-lateral axis showed no significant effect on ECAP latency. (H) ECAP magnitudes showed variable subject-specific changes with medial-lateral lead migration.

To investigate the effect of medio-lateral shift in the stimulation lead on the ECAP morphology recorded on the second lead, the stimulation lead was shifted approximately 3.5 mm laterally from the midline (Fig 5E). Fig 5F shows ESR traces form a single subject during a shift from medial to lateral – showing a small decrease in latency and a decrease in ECAP component AUC. Data analysis for ECAP characterizations were replicated in the same fashion as data analyzed within lead migration along the rostral-caudal axis. Lateral shift of the stimulation lead showed no significant differences to the ECAP latency from the medial position (0.86 ± 0.077 ms) to the lateral position (0.819 ± 0.104 ms), as shown in Fig 5G. The interquartile range for calculated ECAP latency from both medial and lateral positions was 0.12 ms (25-75%: 0.82-0.94 ms), indicating a very narrow spread within the latency measurements. There was an increase in ECAP component magnitudes, reported as percent changes, after shifting the stimulation lead laterally: N1: 82.8 ± 27.2%; N2: 16.3 ± 59.4%; N3: 57.4 ± 5.7%; N4: 196.4 ± 52.7% (P < 0.001) (Fig 5H).

### EMG signal spilling over into ECAP component in ESR

Here, we report on evoked myogenic artifact observed in our ESR recordings. This furthers our understanding of signal sources in the ESR signal and opens further applications of ESR to EES therapy. Fig 6A shows ESR traces from the recording lead (left in red) and EMG traces from bipolar electrodes inserted into the local back muscles (right in blue) from a single subject at several stimulation amplitudes. At 1.2 mA, a distinguishable ECAP component was seen with no EMG component present in the ESR. At 2 mA, an EMG recordings collected from the muscles was apparent with the median trace showing EMG signal with a small amplitude, from about −30 to 20 ŪV peak-to-peak (Fig 6B). At a stimulation amplitude of 3 mA, we observed an ECAP component in ESR with a peak-to-peak amplitude of approximately −40 to 20 □ V. Stimulation at this level also triggered a clear EMG signal recorded in the muscles. The EMG signal collected from the muscles observed was large with a −500 to 1000 □V peak-to-peak value. The insertion in Fig 6A indicates that the median EMG signal had an amplitude from −100 to 100 □ V, which is much larger than the median EMG signal observed at the stimulation amplitude of 2 mA.

**Figure 6:**
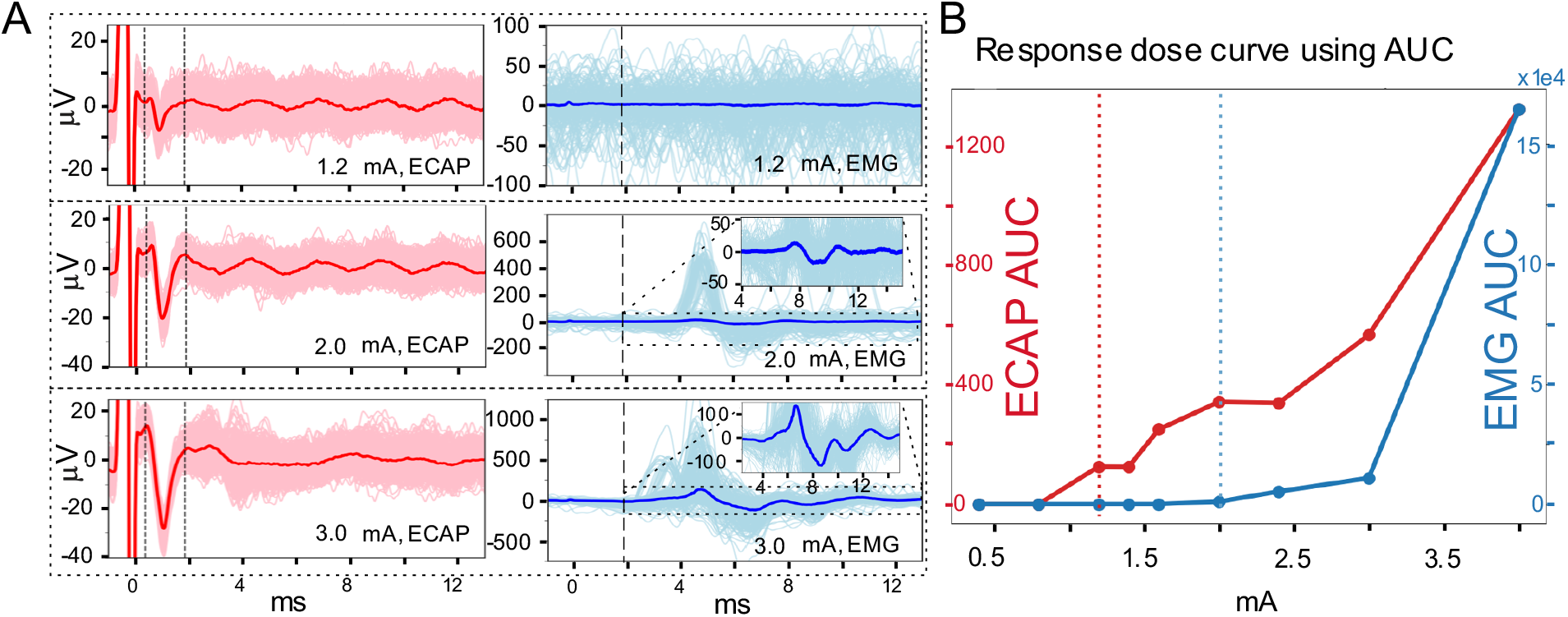
ECAP and EMG dose-response curves. (A) Representative ECAP component in median ESR and EMG recordings (from intercostal muscle between the lowest two ribs) plotted overlain on 300 raw traces for stimulation amplitudes of 1.2 mA, 2 mA, and 3 mA with two leads placed in series. The left column, in red, shows ESR from channel Ch13. The two dashed lines indicate the ECAP component window from 0.33 to 1.80 ms. The EMG window is from 1.8 to 13 ms. (B) Dose response curve of the ECAP component (red) and EMG (blue) signals from (A). The magnitude of the response is calculated using area under the curve of the signal in the defined ECAP and EMG windows shown in (A), respectively. The vertical dotted lines indicate the ECAP and EMG threshold where the first response is observed in each dose-response curve.

We constructed dose-response curves to investigate the overlap between the activation levels of the ECAP component in ESR and EMG signals collected from the muscles. Fig 6B summarizes the ECAP and EMG AUCs at different stimulation currents as dose-response curves. Both ECAP and EMG signal strength increased as the stimulation intensity was increased and the ECAP threshold was lower (1.2 mA) than the intercostal muscle EMG threshold (2 mA) (Fig 6B). This observation is consistent across different channels for the ECAP recordings within the same subject and across subjects (Table 1). On average, the EMG detection threshold in the muscles (EMG/ECAP row in Table 1) is 56% higher than the ECAP detection threshold measured in ESR.

**Table 1.** ECAP and EMG threshold (mA) comparison across different animal subjects.

|  |  |  |  |  |  |
| --- | --- | --- | --- | --- | --- |
| **Subject** | N1 | N2 | N3 | N4 | N5 |
| **EMG** | 2 | 2.4 | 1.8 | 2.4 | 3 |
| **ECAP** | 1.2 | 1.6 | 0.9 | 2 | 2.1 |
| **EMG/ECAP** | 1.67 | 1.5 | 2 | 1.2 | 1.43 |

Notably, the ESR trace at 3 mA of stimulation shows a signal outside the ECAP component window in the 4-8ms time range that corresponds to the time range in the EMG trace where activity is seen. These data suggests that electrical activity from myogenic contraction in the local back muscles spilled over into the ESR traces recorded on the Octrode^™^ leads.

Intramuscular EMG recordings were collected from several muscle groups: intercostal muscle (IM), paravertebral muscle’s rostral part (PVR), paravertebral muscle’s caudal part (PVC), gluteus (G), quadriceps (Q), bicep femoris (BF). Additional recording with needle electrodes inserted in the intercostal muscles across the skin without dissection (IMS) were also collected. Results shown in Fig 7 demonstrate that IM recorded the largest magnitude responses, while the needle inserted in the muscle through the skin resulted in lower magnitude recordings (IM and IMS recording in Fig 7A). PVR and PVC also had activation at this stimulation amplitude, but the EMG magnitude was much lower compared to that from the intercostal muscle (Fig 7A). For the muscle groups where activation was observed, the activation threshold for each muscle was different (Fig 7B). Activation of the GL, Q, and BF muscles was minimal during stimulation.

**Figure 7:**
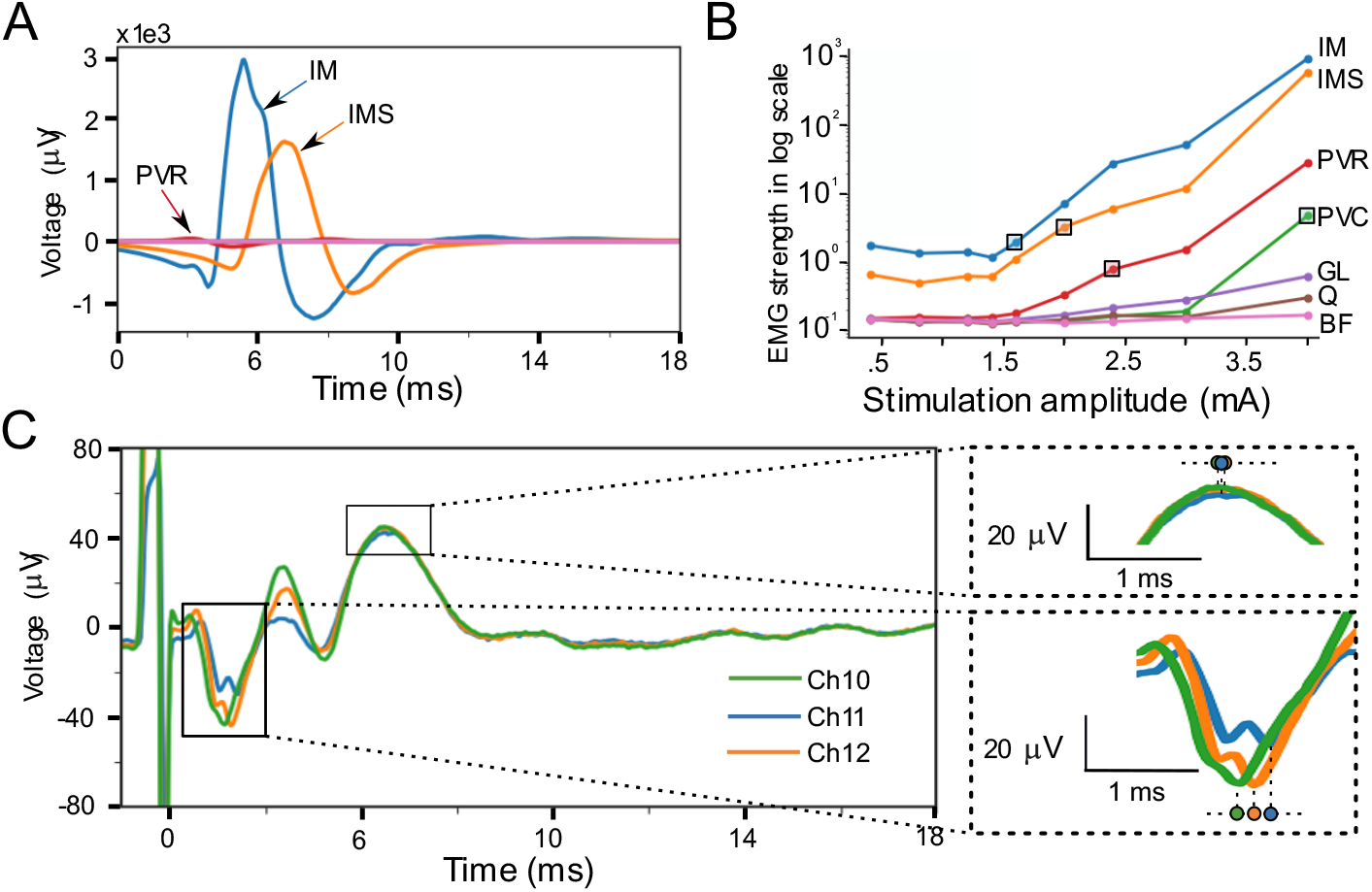
Intramuscular EMG recordings during EES and possible spillover of the EMG signal into the ECAP component in ESR trace. (A) EMG traces recorded by bipolar recording electrodes inserted in or near the muscle group. (B) Dose-response curves of the EMG recordings from bipolar electrodes. Boxes indicate stimulation current at threshold for each muscle activation, among which activation for GL, Q and BF muscles were not observed. The EMG dose-response curves were calculated using the EMG window from 1.8 to 13 ms and the magnitude is displayed in log scale. (C) Representative ESR traces with clear contamination to ECAP component from EMG component spillover. ECAP component versus EMG component spillover in ESR can be differentiated as the ECAP has a known conduction velocity that results in a signal propagation delay across recording contacts. Insertion on the right showing signal propagation delay in ECAP component but not EMG component. Each recording trace corresponds to a median waveform of 300 trials and a stimulation amplitude of 4 mA.

Evoked muscle activity was also recorded by the contacts on the Octrode^™^ lead, as shown in Fig 7C, where the EMG component was present alongside the ECAP component in ESR and possibly overlapping with ECAP component. It is important to be aware of the various artifacts that can appear in ESR recordings, which can be mistaken as ECAP component. To differentiate authentic ECAP component from EMG bleed through, due to activation of nearby muscles, we compared the signal propagation delay across recording contacts on the lead. An ECAP propagates at a finite speed generally in the range of 30-70 m/s (for A*β* fibers in dorsal column) and consequently appears at a delay of ~0.1 ms between adjacent contacts, while an EMG signal appears simultaneously on adjacent recording contacts (Fig 7C breakout). Differentiation of ECAP vs. EMG is illustrated in blow outs on the right side of Fig 7C. Contamination of the ECAP component with EMG artifact in ESR could complicate ECAP-only oriented analysis and closed-loop control algorithms implemented in an implantable device.

### Effect of reference electrode location on morphology of EMG artifact

Fig 8A shows that the recording channel chosen can have a strong effect on the morphology and magnitude of the EMG artifact recorded during ESR. These plots show that as the recording contact is moved caudally closer to the reference electrode contact and further from the stimulation contacts the amplitude of the EMG artifact decreases while the latency only minimally increases. The DIFF-V plot has a notable overall reduction in EMG size when compared to the other plots. The effect of referencing on EMG is seen in Fig 8B with DIFF-V having a dramatically reduction in overall amplitude of EMG artifact. REF9-V has a similar EMG shape to LTR but is slightly shifted so that the peak comes earlier and the max and min is lower than when using LTR. The peak also seems to appear earlier leading to the conclusion that as the reference electrode is moved closer to the recording electrode the latency between the stimulation artifact and the EMG artifact decreases. In Fig 8C the AUC of the EMG artifact was measured for the time window from 3.5-9ms. The EMG component AUC was significantly higher with LTR (normalized to 1 in each of n=5 subjects) when compared to the DIFF-V referenced recordings (0.278 ± 0.123 SE) (P<0.005 for DIFF compared to LTR) (Fig 8D). There was no significant difference seen between the LTR (normalized to 1 in each of n=5 subjects) when compared to REF9-V referenced recordings (.965 ± 0.287). The variability in REF9-V could be a result of variation in LTR reference electrode placement across subjects

**Figure 8.**
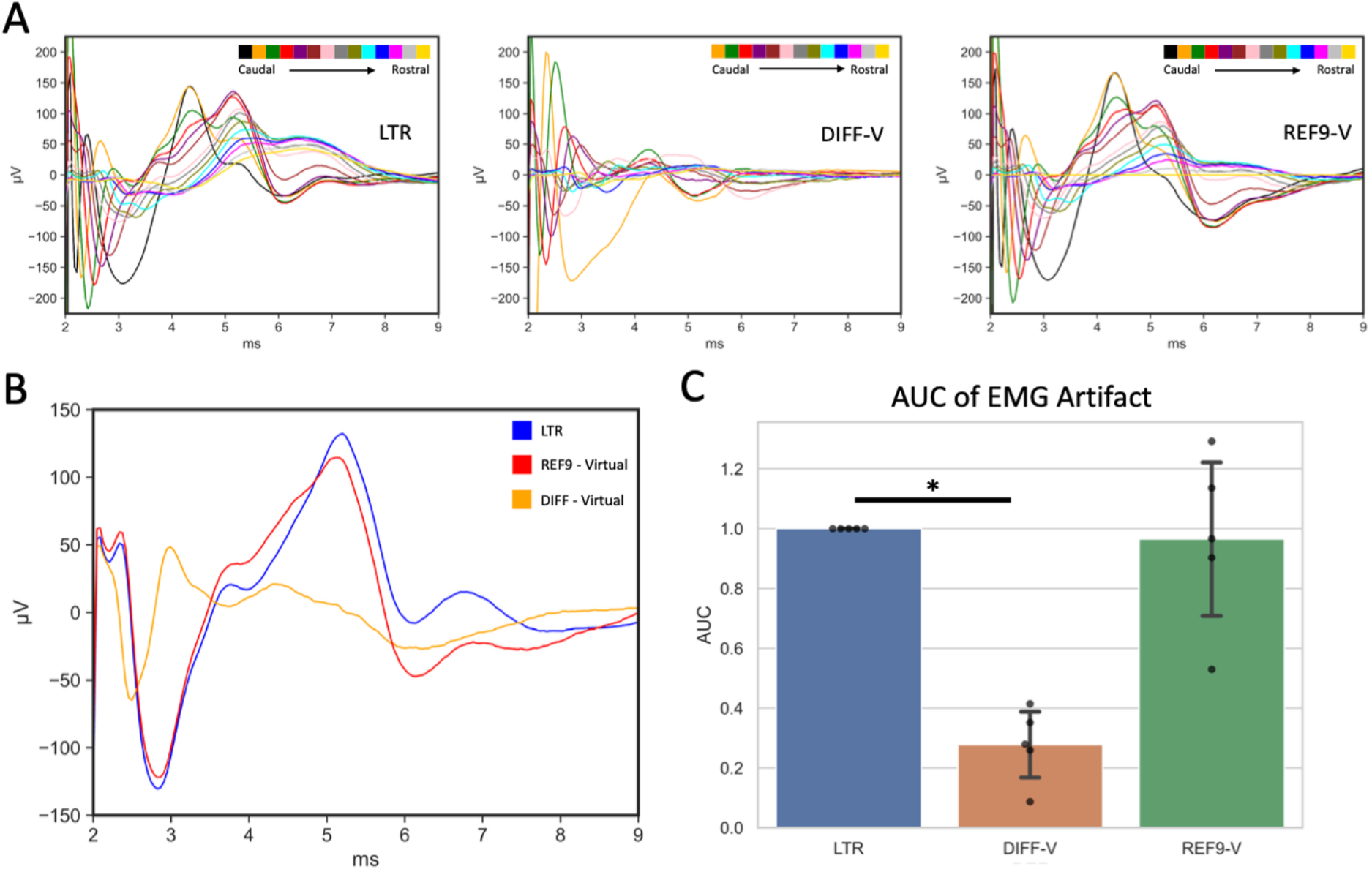
Effect of referencing on EMG artifact morphology and latency: A) Representative recordings on all available recording channels demonstrate changes in EMG magnitude and morphology with local tissue reference (LTR), virtual differential reference (DIFF-V), virtual contact 9 as reference (REF9-V) and contact 9 as reference (REF9). Each recording trace is a median waveform of 150 trials for a stimulation amplitude of 4.5 mA. B) Representative plot of EMG artifact recorded on channel 1 (Ch1) using different referencing options: local tissue reference (LTR), virtual differential recording reference using channel 2 and recording from channel 1 (DIFF-V), and virtually referencing contact 9 (REF9-V) at a simulation amplitude of 4.5mA C) AUC (Absolute Value Sum) of EMG artifact using 3.5-9ms time window post stimulation pulse. * Indicates significant difference in the ECAP magnitude (p<0.05).

## Discussions

The ESR collected from the implanted leads in the epidural space captures the stimulation evoked neural and myogenic response of tissue around the recording electrode during EES. Existing work has shown the utility of the ECAP component in continuously adjusting stimulation amplitude to prevent overstimulation and an excessive sense of paresthesia to the patient. The ECAP component in ESR may provide further clinical utility. However, it is inappropriate to mistake the entire ESR as ECAP signal, as ESR will contain other physiological and non-physiological information.

In this study, we first reviewed the fundamentals of electrophysiology recording and particularly the role of referencing strategy on stimulation artifact and its impact on ECAP morphology in ESR. Secondly, we investigated ECAP features to detect lead migration, a common adverse event in EES. Lastly, we presented and characterized the myogenic response present in certain ESR recordings to initiate the exploration of applications of myogenic responses in ESR. We used a swine animal model in this study because previous studies on the effect of EES in swine have demonstrated similar result to those found in the human sensorimotor system, suggesting that the porcine model is a relevant translational model to humans (R. Islamov et al., 2020) (R. Islamov et al., 2022)(R. R. Islamov et al., 2017)(Fadeev et al., 2020).

### ECAP component and stimulation artifact morphology in ESR is affected by referencing strategy

One factor affecting ESR that has not been well investigated is the referencing strategy. When recording ESR during EES, there are multiple ways that one can reference the recordings. In this study, we investigated the following reference strategies: reference placed in nearby nonexcitable tissue (LTR), differential reference of two neighboring channels (DIFF), and reference to one of the contacts on the lead. The results showed that LTR provided the best recordings of ESR, with consistent ECAP component latency between contacts, consistent morphology, and large signal amplitude compared to other referencing methods (Fig 2) (Fig 3). Using LTR based recordings to estimate the conduction velocity of ECAPs fell into the range consistent with previous works. LTR also provides flexibility of referencing location (Fig 1). Clinically, LTR may be implemented using the case of the implantable pulse generator as the reference electrode. In comparison, DIFF recordings had lower amplitude stimulation artifact and ECAP component magnitude, but also resulted in less channels that can be recorded from (Fig 2) (Fig 3). On the other hand, using contact 9 or 3 as reference substantially changed the morphology of ECAP component with REF3 causing obvious distortion in the ECAP morphology and conduction velocity estimation (Fig 2)(Fig 3). REF3 also resulted in a flipping of polarity at contact 3 due to the relationship to the stimulation dipole changing as the recording contact moved above contact 3. One reason for the change in ESR morphology is that the stimulation artifact heavily contaminated the recorded signals when using contact 3 on the lead as the reference electrode (REF3). This was due to the proximity of REF3 to the stimulation site. REF9 also suffered from small morphology changes to the ECAP component but not from the stimulation artifact. When using REF9, on the rising phase of the N1 peak there is a consistent dip in the ECAP component, which is a result of the reference electrode recording the ECAP at the same time as the recording electrode (Fig 2B) and the two recordings ‘subtracting’ together. Despite the change in morphology REF3 and 9 have the advantage of using only existing electrodes, and with proper signal analysis they have the potential to be used as referencing methods for ESR during EES. In this study, we found that referencing plays a critical role in ESR and its interpretation.

### Movement of leads significantly affected ESR morphology

Daily activities of patients include walking, sitting, standing, twisting, and bending over. These actions can all lead to small temporary movement of the leads in the epidural space resulting in suboptimal stimulation. In addition, these movements can cause a permanent lead migration, leading to loss of efficacy in EES therapy (Dombovy-Johnson et al., 2022a). Here, we systematically investigated the effect of rostro-caudal and medio-lateral movements in both the stimulation and non-stimulation leads on the ESR. Slight rostral directional movement of the non-stimulation lead resulted in a significant reduction in ECAP component magnitude and increase in ECAP component latency (Fig 4). Even a small 2-3 mm movement caused a decrease in ECAP AUC of ~24% (Fig 4, position I to II). A similar effect was observed when the stimulation lead was moved along the rostral-caudal axis (Fig 5). This observation is consistent with previous studies showing that ECAP amplitude decreased, and latency increased as the ECAP propagated away from the stimulation site (Chakravarthy et al. 2020). Another key reason for the attenuated ECAP component amplitudes in ESR is that the stimulation artifact in ESR attenuates at distances further from the stimulation site (Plonsey & Barr, 1995) and the stimulation artifact can skew the quantification of the ECAP component, as both can be recorded in the same ESR. Therefore, the quantified ECAP component magnitudes in ESR might not reflect an authentic trend of ECAP signals and using such quantification in a closed-loop control system to adjust stimulation could be misleading.

Unlike other animal subjects, for subject N3, we observed a slight increase in the ECAP component magnitude with a caudal movement of the stimulation lead (Fig 5D). A possible reason for this could be that the anatomy near the vertebrate disc makes the stimulation less effective. The movement of the electrode from the more rostral to more caudal position changed the relation of the electrode to the surrounding spinal discs (Supplementary Fig 1). Another possible factor could be that the stimulation electrode might have moved from a position that is far away or off a dorsal rootlet to a position that is closer or on the rootlet. As a results, there is a larger chance that the rootlets were activated, which in turn activated nearby muscles in the back, contaminating the ECAP component in ESR. The duration of the negative peak in the ECAP component from subject N3 was wider when stimulation was in the caudal position, while that response has a narrower negative peak when the lead was placed in a more rostral position (Supplementary Fig 1), supporting the presence of the longer duration EMG component into the ECAP component in the same ESR. These reasons may explain why the AUC quantification of ECAP component magnitude increased in subject N3 with caudal movement of the stimulation lead. Further work is needed to understand how underlying neuroanatomy and surrounding tissue properties shape ECAP morphology(Anaya et al., 2020) (Parker et al., 2020).

When moving the stimulation lead along the medial-lateral axis, a 2-3 mm lateral movement caused significant change in the ECAP amplitude but not in its latency (Fig 5). As the investigated recording electrode (Ch11) was far from the stimulation site, the impact of stimulation artifact changes were negligible. The results suggest that even a small movement along the medial-lateral axis could cause activation of distinct neural substrates resulting in different ECAP component amplitudes with a similar conduction distance from the evoked responses’ initiation site to the recording site. Notably, the change in the ECAP magnitude was not consistent across all subjects. Lateral movement results in an increase in ECAP magnitude in all subjects except for N3, where the ECAP magnitude decreased. (Fig 5H). A possible reason for this is that the root and rootlets distribution on the dorsal spinal cord may have some animal to animal difference and on N3 the stimulation lead was likely located between the roots. Thus, our, previous studies suggest that variation in signals could depend on position of the electrodes in relation to portions of the dorsal roots or intervals between the roots (Cuellar et al., 2017) (Mendez et al., 2021). A similar movement along the medial-lateral axis could place the stimulation contacts in a very different anatomical locations in different subjects, which may result in activation of nearby muscles through rootlet activation and in turn produce a distinct magnitude change in the recorded ESR. In some cases, the lead might have been also moved with a slight caudal-rostral shift, instead of a pure medial-lateral shift, causing the differing trends observed (Supplementary Fig 1). These results together indicate that ECAP components may be used to analyze the potential lead migration in patients, while further decomposition of the effect from stimulation artifact and EMG signals on the quantified ECAP component in the same ESR should be further investigated. The sensitivity of the ESR to lead position and possibly to the underlying anatomy (e.g., presence of dorsal roots), may also enable an intraoperative and programming application for evoked response sensing to guide lead placement and stimulation contact selection respectively.

### Local back muscle activation during EES therapy

Another interesting observation is that the evoked EMG activity could be recorded in the same ESR where the ECAP component was recorded. This indicates that evoked responses triggered by EES could have both ECAP and EMG components, which could be caused by different neural substrates. The EMG components in ESR likely originate from more than one activated muscle group (Fig 7). A naïve quantification treating the entire ESR from a contact on the ESS lead as the ECAP signal is inappropriate and could miss-lead the therapy adjustment in a ECAP signal based feedback control system. Instead, signal processing algorithms should be developed to carefully decompose these different physiological components before any quantification is applied.

EMG features in the ESR recording could be used to inform optimal stimulation contact selection to engage on-target neural fibers and avoid activation of therapy dose limiting off-target neural pathways. For this application, it is critical to recognize that referencing strategy has a profound effect on the EMG components in ESR. Out data shows that the DIFF referencing resulted in the smallest magnitude EMG components recorded while the LTR and REF9 resulted in larger EMG magnitude components (Fig. x). This is due to the EMG behaving as a common mode artifact, similar to stimulation artifact. Further, we show that EMG across the recording channels can show an apparent conduction delay, which is especially pronounced in the LTR referencing scheme (Fig. x). This apparent conduction delay is produced as there are likely several sources of EMG close to the recording and/or reference electrode along with ECAP components in the ESR. These sources of signal add to create an apparent conduction delay in the EMG components of the ESR. Our study shows that the ESR can contain EMG components. We propose these EMG features may be used to inform therapy delivery and show that referencing strategies affect the EMG feature amplitude and morphology in the ESR.

### General limitations

During this acute large animal study, the partial laminectomies were performed at several levels as described in the Methods. The laminectomy could change the electrical properties of tissue surrounding the stimulation and recording leads, which are known to affect neural stimulation and recording (Anaya et al., 2020). Further, the acute bleeding from the procedure introduced electrically conductive fluid around the leads, likely increasing current spread from stimulation. The C arm used for imaging and placement of leads had to be moved after each image resulting in perspective changes between images used for placement decreasing the accuracy of the electrode position. Between subjects there was also some variation between the lead placements which can be seen in supplemental fig 1. When moving leads rostrally and caudally the movements were not strictly rostro-caudal due to some medial lateral movement being induced. This can create variating in stimulation location and distances between contacts on separate leads. Although we presented preliminary data identifying ESR recordings contain evoked EMG artifacts, we did not conduct experiments with muscle paralytics administered to differentiate all EMG from ECAP recordings. It is therefore possible that additional EMG components were present in the ESR that we did not identify. Durring all experiments the pig was under anesthesia with isoflorine, fetynal, and at times buprenorphine which could effect spinal reflexes and in turn our recordings. We believe the findings of the study are robust despite these limitations as most conclusions were based on relative comparisons between, for example, referencing strategies. Care was also taken to minimize fluid accumulation in the surgical field with the EES leads.

## Conclusions

This study evaluates several key challenges in recording and quantifying evoked responses recorded from EES leads. ESR is the recordings collected in the epidural space outside the spinal cord which can include multi-modality physiological and non-physiological signals, such as stimulation evoked ECAP from spinal cord, evoked EMG from local muscles, stimulation artifact etc. The results demonstrate the impact of different referencing strategies on the ESR morphology, ECAP component latency, and magnitude of ECAP and EMG components and the role of minimal movements of stimulation and recording contacts on changes in the recorded evoked response. These findings will lead to further exploring applications for EES evoked responses to inform the response to lead migration, guide intraoperative lead placement, and therapy programming (e.g., stimulation contact selection and stimulation parameters). The results on the activation of evoked EMG signals on both the EES leads and needle electrodes inserted directly in muscles provide preliminary insights on addressing target engagement during EES for optimal neural substrate activation. Future studies should investigate and isolate the different neural substrates activated during EES. In addition, advanced evoked response signal processing algorithms should be explored to carefully distinguish the components of stimulation artifact, evoked EMG responses, and ECAP components present in the ESR to make stimulation evoked responses a reliable objective measurement for EES based therapies.

## Acknowledgements

The authors thank Simeng (David) Zhang for his contribution during the early phase of this project.

