## Supplemental materials for "Characterization and Applications of Evoked Responses During Epidural Electrical Stimulation"

### Supplementary Material

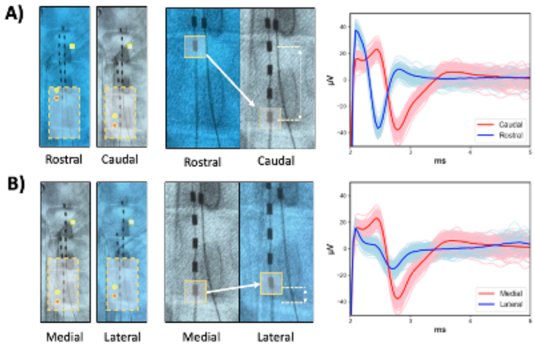


**Supplementary Figure 1:** lead movement for animal subject N3 shown in Fig5. (A) Caudal movement of stimulation lead. (B) Lateral movement of stimulation lead.

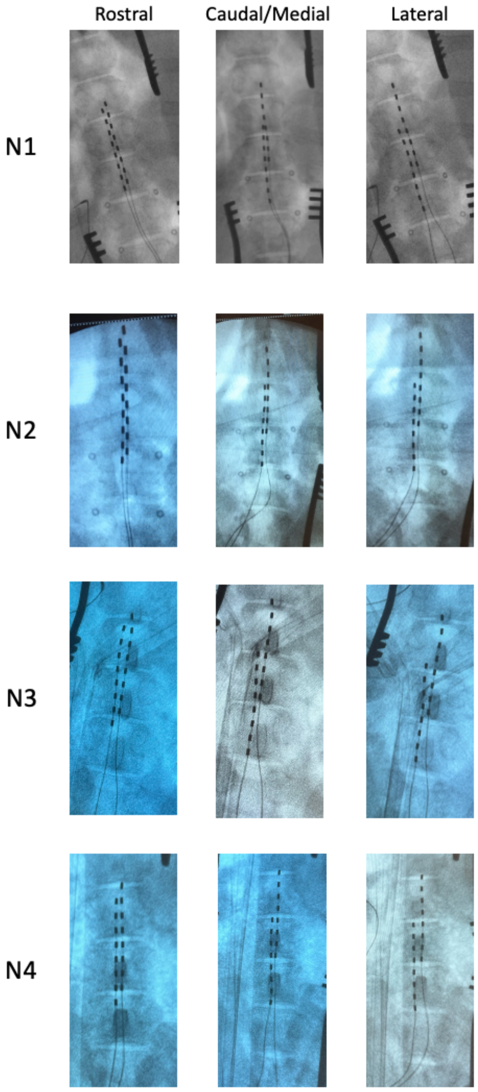


**Supplementary figure 2:** X-Ray images of location used for stimulation.
